# Rational mRNA Structure-guided Attenuation Enables Embryo-safe in-ovo NDV Vaccination and Protection

**DOI:** 10.64898/2026.09.29.755556

**Authors:** Tamir Tuller, Zohar Zafrir, Dana Goldenberg, Avner Finger, Yevgeny Khinitz, Modi Roopin

## Abstract

Newcastle disease virus (NDV) continues to cause major economic losses in poultry, driving many recent virulent outbreaks. Maternally derived antibodies (MDA) can interfere with the development of active immunity following vaccination in young chicks. NDV vaccine strains that are safe for day-old chicks are not safe for administration to embryos.

Here, we combined a computationally guided attenuation strategy with an in-ovo vaccination and challenge study to develop and evaluate an embryo-safe NDV vaccine candidate. Using a large NDV sequence compendium, we applied a local RNA folding energy based algorithm to identify folding selected clusters within the NDV fusion protein; we introduced targeted synonymous substitutions within the first ∼900 nucleotides to modulate local RNA structure while preserving the encoded protein sequence. The recombinant attenuated variant was propagated in embryonated chicken eggs to generate vaccine stocks for in-ovo vaccination. After 18 days of embryonation, the synthetic variants were well tolerated, achieving more than 97% hatchability, with no pre-challenge morbidity or mortality, while unvaccinated challenge controls showed complete mortality.

Together, these data support a model in which structure-aware synonymous modifications can yield an embryo-safe NDV backbone suitable for in-ovo delivery, with protective efficacy exhibiting a clear dose “window” under high MDA conditions.

## Introduction

Newcastle disease (ND) remains one of the most economically important viral diseases of poultry worldwide, despite the extensive implementation of vaccination programs [1–3]. The Newcastle disease virus (NDV) exhibits substantial genetic diversity, and virulent strains belonging to several genotypes continue to circulate in the global poultry populations. For example, Genotype VII have been associated with major outbreaks across multiple geographic regions [4–7]. Although vaccines available at present are generally effective in reducing clinical disease and mortality, genetic and antigenic differences between vaccine strains and circulating viruses may influence protection against infection and virus shedding [4, 7].

NDV strains have traditionally been categorized as velogenic, mesogenic, or lentogenic according to their pathogenicity in chickens. Lentogenic genotype II strains, including La Sota and closely related vaccine strains such as VH, are widely used as live vaccines because of their low pathogenicity and favorable immunogenic properties [8–11]. The commercial VH strain is closely related to La Sota, with greater than 99% amino-acid identity in both the F and HN proteins [11]. The fusion (F) protein is a major determinant of NDV pathogenicity and is initially produced as an inactive precursor F0, which requires proteolytic cleavage into F1 and F2 to become fusion competent. Although NDV virulence is a complex multigenic trait, the amino-acid sequence of the fusion cleavage site (CS) represents a major molecular determinant of pathogenicity and tissue tropism. Avirulent strains typically contain a monobasic CS motif that can be processed only by a restricted range of host proteases, whereas virulent strains generally contain a multibasic CS that permits activation by broadly distributed intracellular proteases and facilitates systemic spread [8, 9, 12].

Vaccination remains the cornerstone of ND control; however, achieving reliable protection during the early post-hatch period presents specific challenges. Maternally derived antibodies (MDA) provide important passive protection to young chicks but can interfere with replication and immune stimulation by conventional live NDV vaccines administered after hatch, thereby delaying the development of active immunity [13–16]. Hence, in-ovo vaccination offers an attractive alternative by allowing standardized and automated vaccine delivery during embryonic development and initiating immune priming before hatch. This approach may therefore help reduce the period of susceptibility associated with declining maternal immunity [16]. The direct use of conventional live NDV vaccines for in-ovo administration, however, is constrained by the susceptibility of the developing embryo to NDV replication. Live vaccine strains routinely used after hatch may cause embryo mortality, impaired hatchability, or post-hatch morbidity when administered in-ovo [17–19]. Earlier attempts to overcome this limitation included attenuation of La Sota derived viruses through selection of escape mutants, which reduced embryonic pathogenicity while retaining the ability to induce protective immune responses [18]. Nevertheless, the challenge of achieving sufficient embryo safety while maintaining adequate post-hatch vaccine replication and immunogenicity has remained a major obstacle to the development of live NDV vaccines for this route. Several subsequent studies have demonstrated the feasibility of developing live NDV candidates suitable for in-ovo administration. The thermostable TS09-C strain, derived from the V4 strain, showed favorable embryo safety and conferred protection against virulent NDV challenge following in-ovo vaccination of specific-pathogen-free chickens [19]. Subsequent evaluation in commercial chickens carrying MDA demonstrated that TS09-C remained protective despite maternal antibodies, providing at least 80% protection against virulent challenge for up to three months [16]. Recombinant NDV candidates generated using reverse-genetics approaches have also shown improved suitability for in-ovo vaccination. Marcano *et al.* reported that an attenuated recombinant NDV candidate improved hatchability and post-hatch survival while maintaining strong protection against virulent challenge [20]. Dimitrov *et al.* further demonstrated that recombinant NDV-based in-ovo vaccines could overcome MDA interference, induce early humoral immune responses, and provide high or complete protection against virulent challenge as early as two weeks after hatch [21].

Studies examining the molecular determinants of embryo attenuation have further highlighted the importance of the F protein. Feng *et al.* demonstrated that the F-protein CS was a critical determinant of attenuation of TS09-C in chicken embryos, supporting the F protein as an important target for rational modification of NDV intended for in-ovo use [22]. However, attenuation strategies do not invariably produce an embryo-safe phenotype. Codon pair de-optimization of the F and HN genes of La Sota, for example, did not provide sufficient attenuation for successful in-ovo application, with substantial adverse effects on hatchability and post-hatch survival [23]. These findings emphasize the narrow balance between attenuation sufficient to ensure embryo safety and preservation of viral fitness required for effective immunization. Recent studies demonstrate continued interest in live NDV-based in-ovo vaccination. In 2026, the TS09-C strain was reported to provide protection against virulent NDV for at least 12 months following in-ovo immunization, while also reducing viral replication and tissue lesions after challenge [24]. In the same year, the thermostable genotype II D58 strain, administered in-ovo on embryonic day 18, induced early seroconversion and provided complete clinical protection in the reported experiment against heterologous genotype XIII virulent NDV challenge, together with reduced viral shedding and enhanced cell-mediated immune responses [25]. Collectively, these studies demonstrate the feasibility of developing live NDV vaccines for in-ovo administration while also highlighting the difficulty of achieving sufficient attenuation for embryo safety without compromising viral fitness, immunogenicity, or protective efficacy after hatch. Despite substantial experimental progress, live NDV vaccines have not yet become broadly established for routine commercial in-ovo vaccination, and the development of safe and effective live NDV candidates for this route therefore remains an important objective [19–25].

Classical attenuation approaches often relies on passaging viruses in tissue culture under non-native growth conditions; such passages accumulate adaptive mutations, some of which reduce viral fitness in the original host. However, the exact mechanisms by which these mutations lead to attenuated phenotypes, or their contributions to safety, immunogenicity, and evolutionary stability are not completely understood [26, 27]. Advances in computational biology, genome synthesis, and reverse genetics now enable viral attenuation to be approached as a rational, multi objective, sequence design problem. Synonymous modifications can affect all aspects of gene expression regulation including translation, mRNA stability, co-translational protein folding and more [28–35]. Previous studies suggested that targeted modification of NDV regulatory determinants can impair viral RNA synthesis, transcription, replication, and pathogenicity. Moreover, distinct selective pressures acting on NDV RNA and protein sequences suggest that parts of the genome may encode functionally relevant to RNA level information beyond its mere protein sequence [36–38]. It was shown that computational framework for rational viral attenuation using silent mutations, which alters evolutionarily selected local RNA structure patterns, can be used for attenuating RNA viruses such as Zika and Hepatitis C viruses [28, 34, 35]. Building on these principles, the present study applies for the first time rational mRNA structure guided engineering to generate an embryo safe NDV vaccine candidate suitable for in-ovo administration, while retaining the capacity to induce immunity.

Embryonated eggs vaccinated with known lentogenic viruses with dosages higher than 10^2.5^ were previously shown to hatch with sever clinical symptoms, illness, and mortality. Thus, in this study, we deployed our generic computational pipeline for generating live attenuated NDV. Our goal was rational design of a strain with reduced viral replication in fertilized chicken embryos while maintaining efficient production performance. To this end, we devised and evaluated a recombinant NDV variant vaccine, called *in ovo vaccine based on recombinant NDV* (IoV-rNDV), designed to enable safe in-ovo delivery and effective early immune priming. Specifically, we wanted to assess the following: (*i*) embryo safety and hatchability of IoV-rNDV following in-ovo administration, the (*ii*) ability of IoV-rNDV to induce humoral responses under strong maternal immunity, and the (*iii*) level of protection conferred against a velogenic NDV challenge.

## Results

### Overview of the Live Attenuated Virus Pipeline

We have previously devised a comprehensive computational pipeline for the rational design of attenuated live viruses [34, 35, 39, 40]. We now devised an enhanced workflow to evaluate a potential vaccine for NDV. Briefly, we conducted a large-scale computational modeling and analysis to location regions in the F protein that demonstrate patterns of selection for strong or weak RNA secondary structure (*i.e.*, the RNA is closed or open, respectively). Following, we rationally designed an attenuated synthetic variant of the virus, based on these regions, by introduction of silent mutations affecting those regions; specifically, the changes were designed to weaken regions with strong mRNA folding in the NDV genome, and vice versa (see Methods, Computational Approach and Models section). Next, and after DNA synthesis and genome assembly, the modified IoV-rNDV virus was rescued and validated using Fluorescent Focus Assay (FFA; see Methods, In-Vitro section). Following, the virus was propagated in SPF embryonated chicken eggs and harvested to generate high-titer virus stocks. The IoV-rNDV virus was injected to a new stock of SPF embryonated chicken eggs, which were then returned to incubation until hatch. After hatching, chickens were monitored for their antibody levels using HI assay and challenged with NDV GVII Velogenic wild-type strain 30 days after hatching; the birds were monitored for 14 days post-challenge, and mortality was recorded daily. Finally, the results were analyzed and validated (see Methods, In-Ovo section). The study workflow can be seen in Figure 1 and in supplemental figures SF1 and SF2.

**Figure 1.**
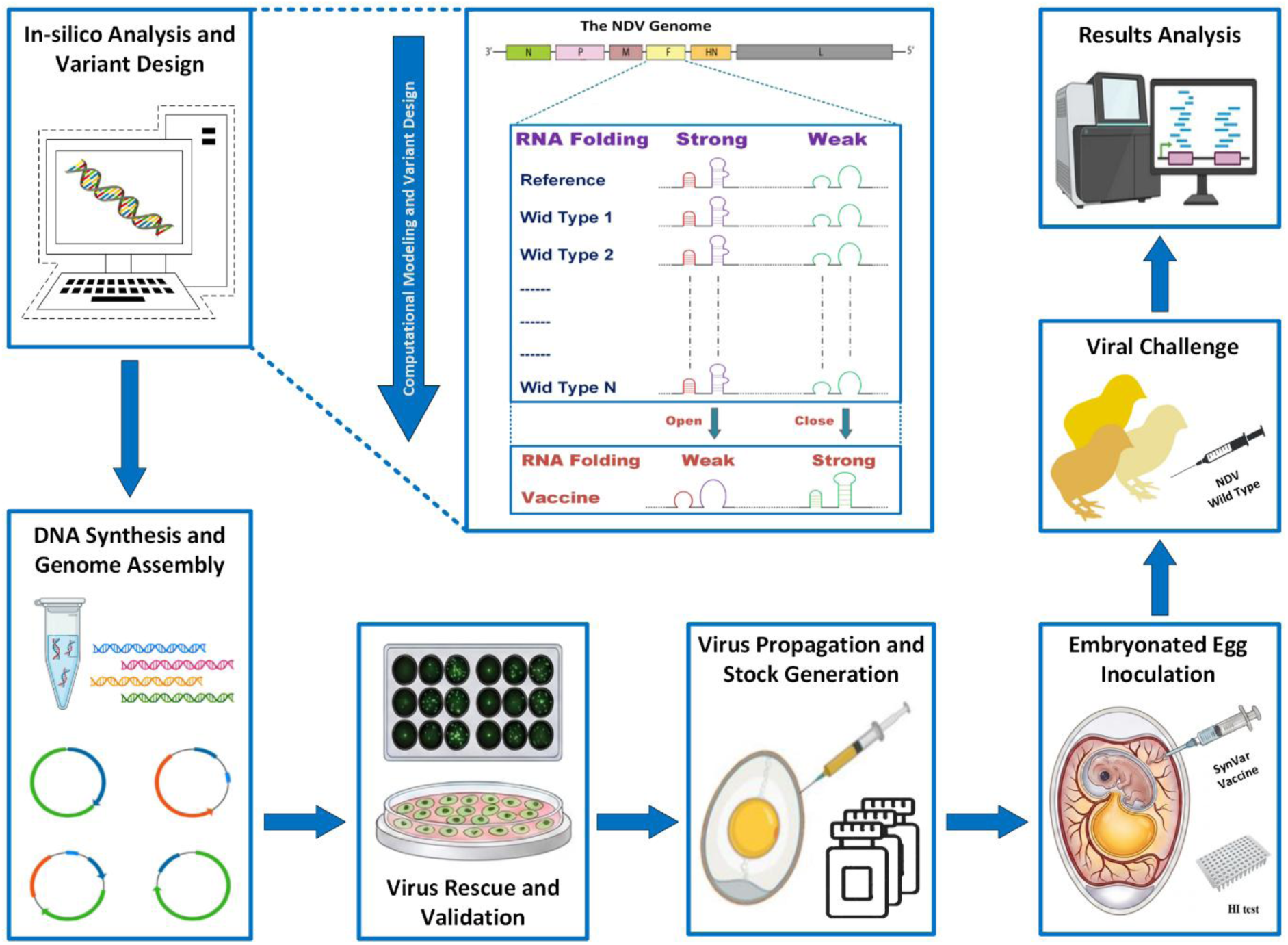
Schematic workflow of the synthetic variant design, rescue, and propagation of Newcastle disease virus (NDV) variants. A) NDV In-silico Sequence Analysis and variant design: A comprehensive NDV sequence database is compiled and analyzed to identify highly conserved RNA secondary folding structures. Synonymous (silent) mutations are introduced, gradually altering the native nucleotide pattern to generate an attenuated version of the virus. B) DNA Synthesize and Genome Assembly: The modified sequence contains a 1,200bp fragment spanning the end of the matrix (M) protein and the fusion (F) protein open reading frame (ORF). The wild-type sequence is replaced with the modified, pattern-altered one to yield the final synthetic NDV genome plasmid. C) Virus Rescue and Validation: the synthetic plasmids co-transfected into BHK-21 cells alongside a T7 polymerase expression helper plasmid and supporting plasmids encoding the viral NP/P/L genes to drive transcription and viral assembly. Successful virus rescue is confirmed via a fluorescent focus assay (FFA). D) Virus Propagation & Stock Generation: Recombinant virus-containing cell lysates are inoculated into 9-day-old SPF embryonated chicken eggs. Following incubation, the virus-rich allantoic fluid (AF) is harvested to generate high-titer virus stocks. E) Embryonated egg Inoculation: 175 broiler eggs are separated into groups, inoculated with the synthetic vaccine, and returned to incubation until hatch. After hatching, HI assay results are recorded at 0, 6, 15, 23, and 29 days of age (DOA). F) Viral Challenge: Birds are challenged at 30 DOA with NDV wild-type strain, monitored for 14 days post-challenge, and mortality is recorded daily. G) Results are analyzed and validated.

### In-silico Analysis and Rational Design of the Live Attenuated Virus strains

The computational pipeline begins with constructing comprehensive databases of viral sequences collected from multiple sources, followed by multi-sequence alignment (MSA) and statistical analyses to identify conserved regions based on both sequence and mRNA folding energy, and focusing on the F protein. These conserved regions are then modified through the introduction of silent (synonymous) mutations that gradually alter the conserved patterns while preserving the original protein sequence (see the Methods section). We subsequently evaluated the viral properties of these engineered variants, and introduced additional modifications to the protein sequence that further attenuated the virus; see figure SF1 and Methods (Computational Approach section).

Our objective was to design a specific strain with attenuated viral replication in fertilized chicken embryos without significantly compromising the production process. To achieve this, we applied our previously devised algorithm [34, 41], which detects regions under selection for local mRNA folding or local mRNA folding energy of the mRNA (LFE); this factor is known to influence various aspects of gene expression and viral life cycle. Using LFE analysis, we introduced targeted silent mutations to modulate local structures (*e.g.*, weakening local mRNA folding at specific positions) while preserving other conserved genomic features. Variants were generated using a simulated-annealing based RNA optimizer, ensuring that neighboring modified sites are clustered within regions spanning at least 44 nucleotides; see details Methods and in Figure 2A-B. The variants introduced silent changes within the first 900 nucleotides of the F protein open reading frame (ORF) of the working strain: VH (pNDVH v2020; see also the Methods section and Appendix 1). Specifically, we introduced silent mutations in two clusters (one weak and one strong) with a total change ratio of 30% relative to the original LFE. This resulted in modification of 25 codons and 30 nucleotides in the reference sequence. The resulting synthetic strain is hereafter referred to as the attenuated variant Number 06; detailed information is provided in Figure 2C.

**Figure 2:**
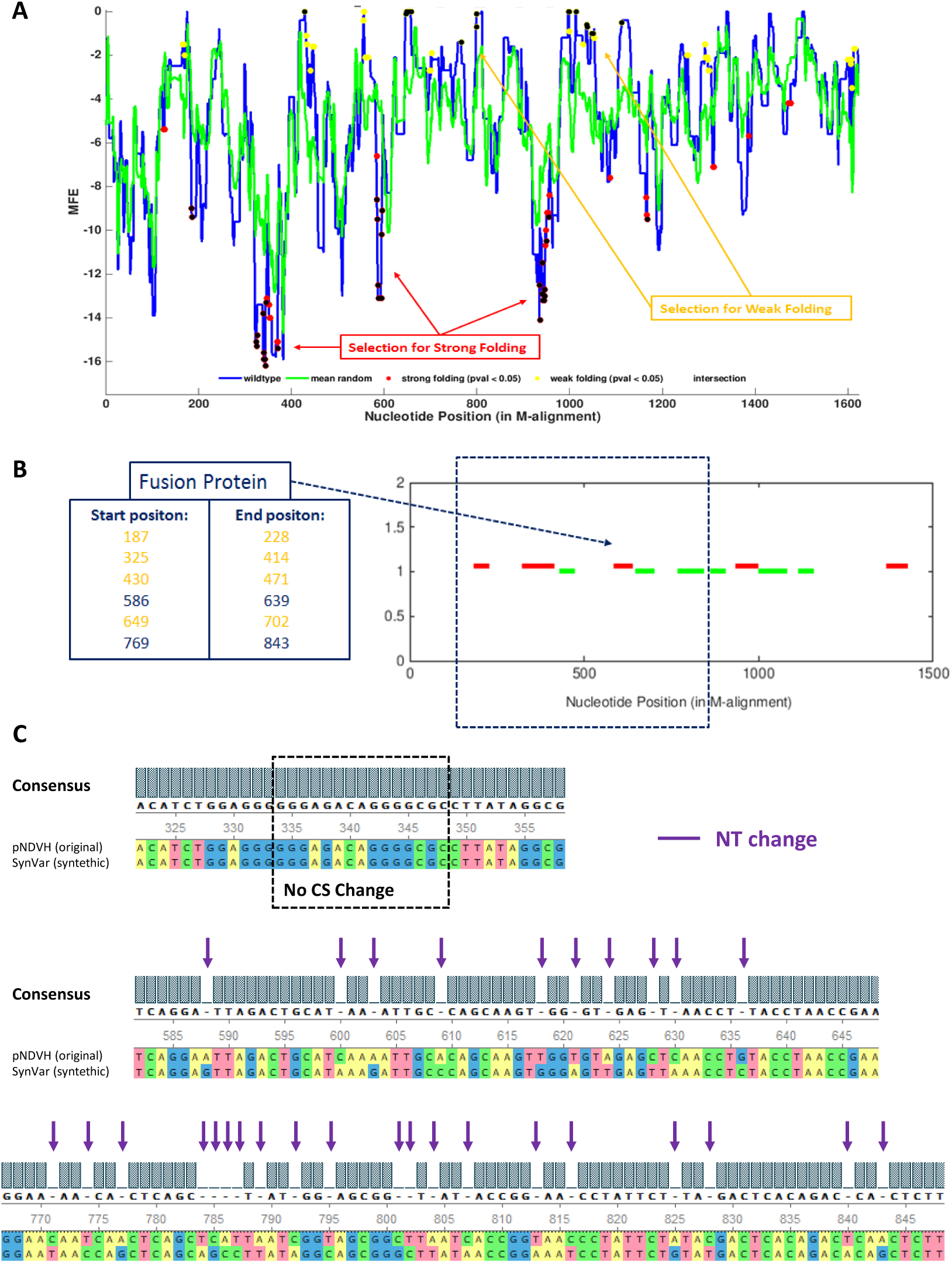
Synthetic variant design stages: Local folding energy (LFE) selection, clustering analysis, and variant modifications. A) Folding selection analysis of the F protein uncovers clusters with significant selection for strong (red) and weak (yellow) folding *(i.e.* low and high LFE, respectively). B) One of three **strong**-folding clusters (red) and one of three **weak**-folding cluster (green) were selected and modified. C) Variant modifications. The Multiple sequence alignment (MSA) of the reference genome and the attenuated synthetic variant 06. The CS (cleavage site) residues at positions 112-117 were **not** modified. The modifications introduced 25 codons, using only silent mutations in 30 nucleotides; the magenta arrows represent silent (synonymous) mutations. The nucleotides alignment shown in the MSA starts at the Start codon.

### Embryonated Egg Inoculation and Viral Challenge

We used three different diluted solutions of 10^1.5^, 10^2.0^, and 10^2.5^ EID₅₀/egg; see details in the Methods, The main safety, serological, and challenge outcomes are summarized graphically in Figure 3. All results can be found in supplementary table ST1-ST3. See also supplemental movies SM1 and SM2

**Figure 3:**
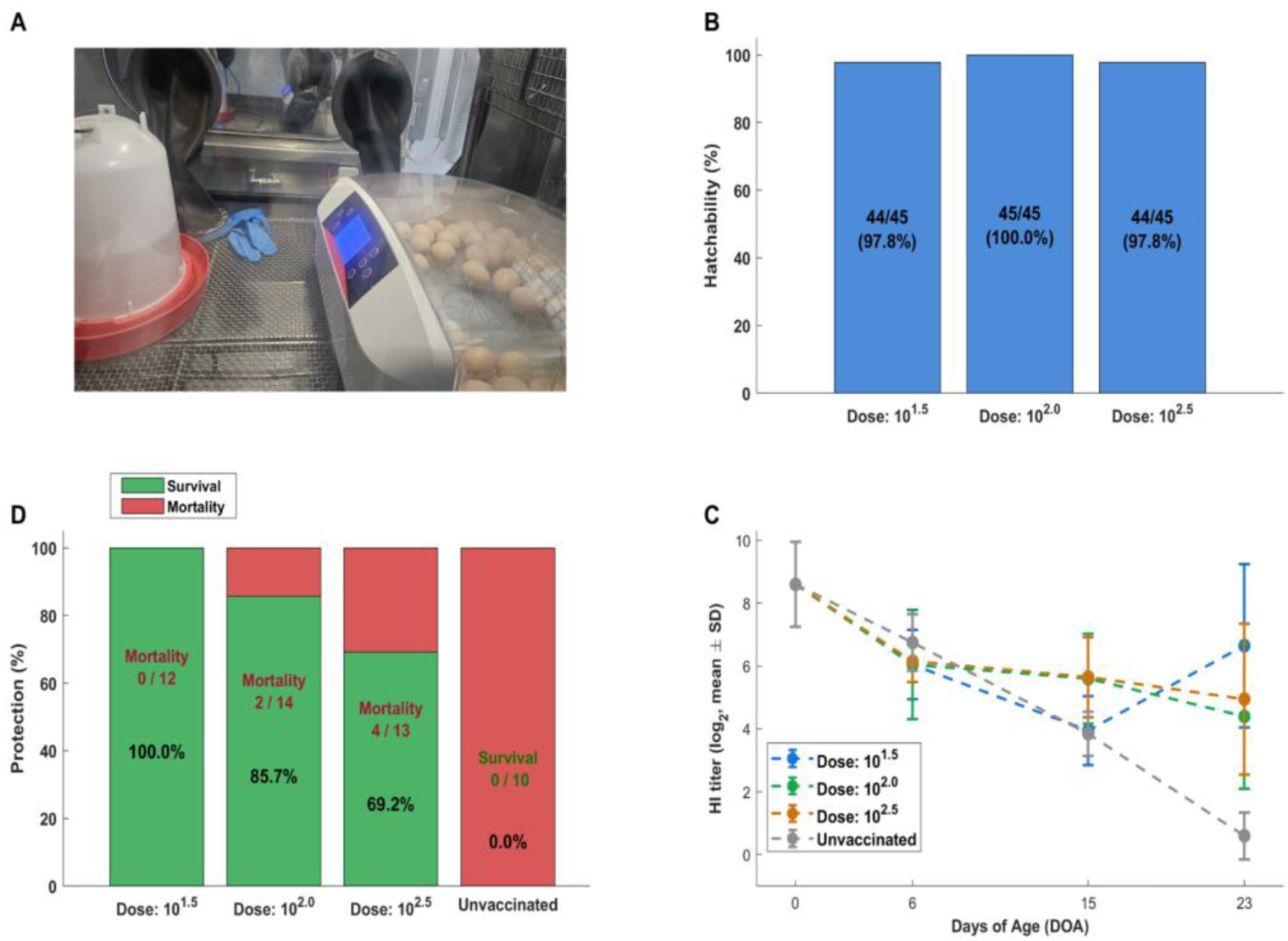
Inoculation, hatching, serological response, and protective efficacy of in-ovo IoV-rNDV vaccination. A) Incubation of the eggs inoculated with the vaccine. B) Hatchability rates in embryos vaccinated in-ovo with IoV-rNDV at 10^1.5^, 10^2.0^, or 10^2.5^ EID₅₀/egg, compared with unvaccinated controls. Values inside bars indicate hatched/total and percentage hatchability. Note that in typical hatching units the typically hatchability with no vaccination is 84–86%. C) HI antibody titers (log₂, mean ± SD) measured at 0, 6, 15, and 23 days of age (DOA) in vaccinated and unvaccinated groups. D) Protection following virulent NDV challenge, shown as survival (green) and mortality (red). Percent survival and mortality counts are indicated within each bar. Hatching picture and additional movies can be found in the supplementary information: supplemental figure SF2, supplemental movie SM1, and supplemental movie SM2.

### Hatchability and Pre-challenge Safety

The inoculated eggs were monitored until hatching (Figure 3A). In-ovo administration of IoV-rNDV was well tolerated at all tested doses (Figure 3B). Hatchability rates were 97.8%, 100.0%, and 97.8% for the 10^1.5^, 10^2.0^, and 10^2.5^ EID₅₀/egg groups, respectively. No clinical signs or mortality were observed in any vaccinated group or in the unvaccinated control group prior to challenge. These findings indicate that in-ovo delivery of IoV-rNDV did not adversely affect embryo viability and showed a favorable pre-challenge safety profile. See details in supplementary table ST1.

### Serological Monitoring

HI titers were monitored longitudinally to evaluate antibody kinetics following in-ovo vaccination (Figure 3C). At hatch (DOA 0), all groups showed identical baseline titers (8.60 ± 1.35 log₂), consistent with maternally derived antibody transfer. By DOA 6, titers had declined in all groups and ranged from mean 6.05 to 6.75 log₂, reflecting the expected early decay of MDA. By DOA 15, divergence between groups became apparent. Mean HI titers were 3.95, 5.60, 5.65, and 3.85 log₂ in the 10^1.5^, 10^2.0^, 10^2.5^, and unvaccinated groups, respectively. The 10^2.0^ and 10^2.5^ groups maintained higher titers at this time point, suggesting earlier vaccine-associated serological activity. By DOA 23, all vaccinated groups remained above the unvaccinated control, with mean titers of 6.65, 4.40, and 4.95 log₂ for the 10^1.5^, 10^2.0^, and 10^2.5^ dose groups, respectively, compared with 0.60 log₂ in the unvaccinated group. Overall, the serological data indicate that vaccination altered the expected maternal antibody decline and supported active humoral responses over time. See more details and standard deviation in supplementary table ST2.

### Protective Efficacy Following Challenge

Protective efficacy following virulent challenge is shown in Figure 3D. Chickens vaccinated in-ovo with IoV-rNDV showed substantial protection against challenge, whereas the unvaccinated control group showed complete mortality. The 10^1.5^ EID₅₀/egg group showed complete protection (100.0%; 0/12 mortality). The 10^2.0^ and 10^2.5^ EID₅₀/egg groups showed 85.7% protection (2/14 mortality) and 69.2% protection (4/13 mortality), respectively. In contrast, the unvaccinated control group showed 0.0% protection (*i.e.* 10/10 mortality). See details in supplementary table ST3.

Taken together, these findings demonstrate that IoV-rNDV conferred strong protection against virulent NDV challenge following in-ovo administration, with the lowest tested dose (10^1.5^ EID₅₀/egg) providing the highest survival.

## Discussion

This study demonstrates that IoV-rNDV, a recombinant NDV variant vaccine, can be safely administered in-ovo, induce measurable early humoral responses in the presence of high maternal antibody levels, and confer significant protection against a velogenic genotype VII challenge at specific intermediate doses. These findings align with growing evidence supporting the feasibility of engineered NDV vaccines for in-ovo use [19, 21].

A major requirement for any NDV-based in-ovo vaccine is embryonic safety. In this study, high hatchability across IoV-rNDV-vaccinated groups (97.7–100%) indicates that the vaccine backbone exhibits appropriate attenuation and restricted replication within embryonic tissues. Similar safety outcomes have been reported for NDV constructs with modified fusion CS or reduced virulence through reverse genetics [17, 22]. The absence of pre-challenge morbidity further supports the suitability of IoV-rNDV for hatchery-level implementation.

A central challenge in the immunology of early-life poultry vaccination is maternal antibody interference. MDA can profoundly inhibit viral replication, particularly for conventional live NDV vaccines administered by parenteral or mucosal routes [5–8]. Consistent with this, all groups demonstrated high HI titers at hatch and at 6 days of age. The divergence observed by 15 days where vaccinated birds at 10^1.5^ and 10^2^ EID₅₀ deviated from the typical MDA decay curve suggests that IoV-rNDV was able to replicate sufficiently to drive active humoral responses under MDA pressure. Similar MDA-bypassing effects have been reported for other recombinant NDV variants designed for enhanced stability or controlled replication [19, 21].

The dose - based patterns observed here further support a biological model in which an optimal replication “window” produces the strongest immune outcomes. Both humoral responses and challenge protection were highest at intermediate doses, whereas the highest dose (10^2.5^) yielded reduced immunogenicity and lower protection. Inverted dose-response curves have been previously described for NDV in-ovo vaccination, where excessive inoculum can limit vaccine take due to early antiviral activity or restricted replication niches in the embryo [17, 22]. Thus, the lack of protection in the 10^2.5^ group is consistent with published dose-kinetic dynamics of live NDV vaccines.

Protection following challenge with a genotype VII isolates a predominant virulent genotype globally [3, 4] revealed strong survivability in the 10^1.5^ and 10^2^ dose groups (100% and 86%, respectively). Although HI titers are commonly used as correlates of protection, immunity to NDV is multifactorial, involving humoral, mucosal, and innate components [42, 43]. In-ovo administration exposes the developing embryo’s respiratory and gastrointestinal mucosa to viral antigen, which may contribute to early mucosal imprinting even in the presence of MDA [44]. While mucosal immunity was not evaluated in this study, the discrepancy between HI titers and survival particularly in the highest dose group supports the idea that non-humoral immunity contributes to protective outcomes.

We can think about two plausible explanations to the fact that optimal vaccination achieved for titer 10^1.5^ and not higher titers. First, the embryos have pre-existing humoral immunity from their mother (*i.e.* MDA); thus, too high dose of a live attenuated virus may trigger an immediate and aggressive local response. The pre-existing antibodies quickly neutralize the vaccine, stopping it from replicating long enough to train broader immune system [15, 45–47]. Second, it is possible that a too high dosage will cause the immune system to overreact locally to the immediate viral load (causing inflammation) rather than building long-lasting memory [48–51].

Overall, these findings identify IoV-rNDV as a promising candidate for in-ovo ND vaccination. The vaccine demonstrated excellent embryo safety, the capacity to elicit early immune priming under high maternal antibody conditions, and meaningful protection against a contemporary velogenic NDV challenge. Further research is warranted to characterize mucosal and innate immune responses, refine dose selection, and evaluate field applicability under commercial production settings. The combination of early-life protection, compatibility with automated in-ovo delivery, and favorable safety profile supports the potential of IoV-rNDV as a next-generation NDV vaccine platform.

## Materials and Methods

### Computational Approach and Models

#### NDV Sequence Database and Multiple Alignment

We generated a comprehensive NDV sequence dataset by compiling sequences from the ViPR (Virus Pathogen Resource) database [52], NCBI GenBank [53], and Wang *et al.* [54]. After filtering duplications and erroneous strains, the final dataset consisted of 623 complete viral sequences. Additionally, we had 2033 full and 4264 partial sequences of the Fusion (F) protein region, respectively. Furthermore, we obtained F protein sequences of the NDV VH strain and Avian paramyxovirus 1 Genotype 7 isolate (GenBank Accession Numbers KC844235.1 and PP928314.1, respectively [53]). The sequences were then aligned, based on their protein sequence, using the Clustal Omega multiple sequence alignment tool (MSA; [55, 56]. The AA-level results were then mapped back to the sequences’ nucleotide level.

#### Null Models

To evaluate the hypotheses of evolutionary conservation and selection for RNA secondary structures, we employed various null models. Choosing various synonymous codons can affect the mRNA folding and thus affect gene expression properties such as translation-initiation, translation-elongations, and protein folding [28, 41, 57–63]. Therefore, we defined two null models: (*i*) Vertical codon usage bias (VCUB) that performs synonymous codon permutations, per column in the MSA analysis; this preserved the MSA codon frequencies and AA for each column independently. (*ii*) Column shuffling (COL_PERM) swaps synonymous codons in randomly chosen columns in the MSA (*i.e.* performs permutation of columns, while only synonymous codons are replaced); this preserves the evolutionary distance between the genes/sequences in the MSA. Specifically, we generate 1,000 randomization models of each sequence in the database based on our random models, *e.g.* for each sequence we generated 1,000 COL_PERM-based models. Following, the actual measure/score at each position was then compared to the set of values received in that position for the 1,000 random alignments and a Z-score value was calculated. See more details here: [28, 29, 34].

#### Variant Design Based on RNA Secondary Structure Analysis

The designed variants are based on our algorithms for detecting selection for local mRNA folding of the mRNA (or local folding energy; LFE), which is known to be able to affect various aspects of gene expression, fitness, and viral life cycle; see some examples in [28, 34, 41, 57]. We only considered positions with a significant selection for high / low local LFE (both VCUB and COL_PERM null models; p_1_, p_2_ < 0.05; Z-score > 2, Z-score < -2), with cluster neighboring locations of at least 44 nucleotides.

Based on the LFE selection analyses we designed a variant, which introduce silent (or synonymous) mutations to modulate such local structures (*e.g.*, weaken positions with selection to strong local mRNA folding) without affecting other conserved features in the viral genome. The modification were introduced to the coding sequences alone, excluding the CS; see figure 2C, and the supplementary information. To this end, we used RNA optimization program (simulated annealing based) to introduce changes in the LFE relative to the original ones. Next, using the modified RNA de-optimizer program, which replaces synonymous codons but does not use rare ones (*i.e.* codons with frequency of less than 15%), we generated gradient of changes to the local folding energy at positions that were detected to be functional based on the selection analysis.

### In-Vitro

#### Production of the Variant Virus

The NDV variant genome was generated by replacing a 1,200-bp fragment spanning the Matrix (M) and Fusion (F) protein interspace junction in the NDV VH genome with a modified sequence. The modified fragment was synthesized *de novo* and cloned into the original genome by Gene Universal Ltd service platform. The final construct was verified by Sanger sequencing (Gene Universal Ltd). Lyophilized plasmid stock, provided at a concentration of 5 µM by Gene Universal Ltd, was re-suspended in ddH₂O and stored at −20°C until use.

#### Preparation of Variant NDV Genome DNA for Virus Rescue

The NDV VH plasmid genome was transformed into *E. coli* NEB10β competent cells (New England Biolabs). Transformed cells were grown overnight on agar plates containing 50μg/mL kanamycin. To confirm the presence of the modified genome, randomly selected colonies were screened via colony PCR using gene-specific primers, and sequencing was performed to verify the integrity of the cloned fragment. Verified clones were further expanded in *E. coli* by overnight growth at 37°C in 200 mL of LB medium supplemented with 50μg/mL kanamycin. Cells were harvested by centrifugation at 5,000 × g for 10 min, and plasmid DNA was purified using the ZymoPURE™ II Plasmid Prep Kit (Zymo Research) according to the manufacturer’s instructions. The resulting purified DNA was used for virus rescue.

#### Virus Rescue and Propagation

Virus rescue was performed by co-transfecting BHK-21 cells (ATCC, CCL-10) with a DNA plasmid mixture comprising of pCAGGS/T7, the variant NDV VH genome plasmid, and supporting plasmids encoding the NP, P, and L genes of the La Sota strain (cloned into pCIneo) at a ratio of 2:2:2:1:1; this is previously described in [8], Briefly, BHK-21 cells were grown to 80% confluency in six-well plates using Eagle’s Minimum Essential Medium (EMEM; Gibco) supplemented with 10% fetal bovine serum (FBS), 1% penicillin/streptomycin, and 1% nonessential amino acids at 37°C under 5% CO₂. Cells were washed twice with Opti-MEM (Invitrogen) prior to transfection with a total of 5 μg of the plasmid mixture using Lipofectamine™ 3000 (Thermo Fisher Scientific), according to the manufacturer’s instructions. Six hours post-transfection, cells were washed with PBS, and the Opti-MEM was replaced with 1 mL of 2% medium containing acetylated trypsin (1.5 μg/mL). At 72 h post-transfection, cells and supernatants were harvested after three freeze-thaw cycles and clarified by centrifugation. Virus rescue was confirmed using a Fluorescent Focus Assay (FFA). For virus propagation, 200–300 μL of transfected cell lysates were inoculated into the allantoic cavity of 9-day-old SPF embryonated chicken eggs, followed by incubation at 37°C for 3 days. Allantoic fluid (AF) was harvested, and virus titers were determined using FFA. AF containing virus was filtered through a 0.22 μm Nalgene syringe filter (Thermo Fisher Scientific), aliquoted, and stored at −80°C as virus stock.

#### Fluorescent Focus Assay (FFA)

96-well plates were seeded with DF-1 cells (ATCC, CRL-3586) at a density of 3x10^4^ cells per well and grown overnight using Dulbecco’s modified Eagle medium (DMEM) supplemented with 2% fetal bovine serum (FBS) at 37°C in a 5% CO_2_ environment, to establish a confluent monolayer of approximately 80-90%. The following day, cells were inoculated with 100µL of 10-fold serially diluted virus samples, with each dilution applied in duplicate. Plates were then incubated for 6 hours (± 0.5 hours) at 37°C under 5% CO_2_ to facilitate virus adsorption. Post-incubation, the media containing the virus was discarded, and cells were washed with Phosphate Buffered Saline (PBS) prior to adding fresh medium and returning the cells to incubation under 5% CO_2_ at 37°C. At 72 hours post-infection, the medium was discarded, and cells were washed with PBS before fixation with 80% ice-cold acetone for 30 minutes. Subsequently, cells were subjected to three washes with PBS containing 0.1% Tween 20 (PBST).

For immunofluorescence staining, cells were incubated with chicken polyclonal anti-NDV primary antibody (Abcam, ab34402; 1:5000 dilution in blocking solution: 1% BSA, 22.52 mg/mL glycine in PBST) for 60 min at room temperature (RT). After washing three times with PBST, cells were incubated with Alexa Fluor 488-conjugated goat anti-chicken IgG secondary antibody (Thermo Fisher Scientific, A11039; 1:500 dilution) for 60 min at RT. To assess monolayer integrity, propidium iodide (Sigma) was added, followed by three PBST washes. Fluorescent foci were visualized and imaged using an IX51 inverted fluorescent microscope (Olympus) at 10× magnification. Foci were manually counted, ensuring that counted dilutions yielded 10–100 foci per well for reliable quantification.

### In-Ovo

#### Viruses and Vaccine Preparation

A modified Newcastle disease virus (NDV) variant vaccine (IoV-rNDV) was used in this study. IoV-rNDV is based on an attenuated NDV backbone engineered for embryo safety and early immunogenicity. Viral working stocks were propagated in 9-to 11-day-old SPF embryonated chicken eggs (ECE). Titers were determined as Embryo Infectious Dose 50% (EID₅₀) following standard procedures. For in-ovo vaccination, vaccine stocks were diluted in brain–heart infusion (BHI) medium supplemented with antibiotics (gentamicin, penicillin, amphotericin B) to prepare inoculant containing 10^1.5^, 10^2^, or 10^2.5^ EID₅₀ per 0.1 mL. Diluted working stocks were back-titrated in SPF ECEs to confirm actual titers. An uninoculated negative control group received 0.1 mL PBS per egg. The velogenic challenge virus was NDV isolate 358, belonging to genotype VII, obtained from the Kimron Veterinary Institute (IVS, Israel). The virus was propagated and titrated in SPF ECEs and used at 10³ EID₅₀/mL for challenge.

#### Eggs and Incubation

A total of 175 commercial broiler eggs were obtained from breeder flock 55689, originating from the same hatchery batch. Eggs were incubated at 37.5 °C and 55–60% relative humidity until 18 days of embryonation (DOE). Maternal antibody status of the breeder flock was not quantified prior to the study; baseline anti-NDV antibody levels were assessed at hatch using HI assay.

#### In-ovo Vaccination

At 18 DOE, eggs were randomly assigned into four groups:

IoV-rNDV 10^1.5^ EID₅₀ (n = 45)

IoV-rNDV 10^2^ EID₅₀ (n = 45)

IoV-rNDV 10^2.5^ EID₅₀ (n = 45)

PBS in-ovo negative control (n = 40)

Vaccination was performed manually using sterile technique. Each egg was disinfected at the injection site and inoculated with 0.1 mL vaccine into the amniotic cavity, using 1 mL syringes with 24-gauge, ½-inch needles. Control eggs received 0.1 mL of sterile PBS. Following vaccination, eggs were returned to incubation until hatch.

#### Hatchability and post-hatch Management

Hatchability was recorded for each group at day 0 (21 DOE). After hatching, chicks were transferred to ABSL-2 isolators and housed with ad libitum access to feed and water. Clinical signs were monitored daily. No mortality or NDV-like clinical signs were observed prior to challenge. Blood samples for HI titration were collected at 0, 6, 15, 23, and 29 days of age (DOA). At 14 DOA, flock thinning was performed in all groups to maintain optimal density and welfare conditions.

#### HI assay

NDV-specific antibodies were quantified using the hemagglutination inhibition (HI) assay following WOAH/OIE standard procedures. Serum samples were heat-inactivated at 56 °C for 30 minutes, serially diluted two-fold in V-bottom microtiter plates, and incubated with 4 hemagglutinating units (4 HAU) of NDV antigen. A 0.5–1% suspension of chicken red blood cells (RBCs) was added, and plates were incubated until RBC controls formed a clear button. HI titers were recorded as the reciprocal of the highest serum dilution inhibiting. hemagglutination and expressed as log₂. Titers ≤ 3 log₂ were considered negative. This assay was used to monitor MDA decay and vaccine-associated antibody responses.

#### Challenge Study

Birds were challenged at 30 DOA with velogenic NDV strain 358 (genotype VII). The virus was administered intramuscularly, 0.5 mL per bird, at 10^3^ EID₅₀/mL. Challenge was performed at the Kimron Veterinary Institute (IVS) under approved biosafety procedures. A separate unvaccinated challenge-control group (n = 10) confirmed challenge lethality. Birds were monitored for 14 days post-challenge. Mortality was recorded daily. Protection was defined as survival post-challenge.

Group coding in the IVS report:

Group 1: unvaccinated challenge control, Group 2: IoV-rNDV 10^1.5^, Group 3: IoV-rNDV 10^2^, Group 4: IoV-rNDV 10^2.5^

#### Statistical Analysis

HI titers were analyzed on the log₂ scale. Distributional normality was assessed using D’Agostino–Pearson tests. Because data were non-normal and discrete, Kruskal–Wallis tests were used for between-group comparisons, followed by Mann–Whitney U pairwise tests with Holm correction for multiple comparisons. Within-group changes between 6 → 15 DOA were evaluated by Wilcoxon signed-rank tests (paired). Effect sizes were calculated as rank-biserial correlation (paired) and Cliff’s delta (between groups). Challenge outcomes were compared using Fisher’s exact test. Mortality risk ratios (RR) with 95% confidence intervals were calculated using the Haldane–Anscombe correction when zero counts were present. Wilson 95% CIs were calculated for survival proportions. Statistical significance was defined as p < 0.05.

## Ethics

All animal procedures were performed in accordance with institutional regulations for animal experimentation and were approved by the relevant ethics committee (approval no. NPC-Ab-IL-2308-485-2).

