## Supplementary information for "Rational mRNA Structure-guided Attenuation Enables Embryo-safe in-ovo NDV Vaccination and Protection"

### Supplemental Figures

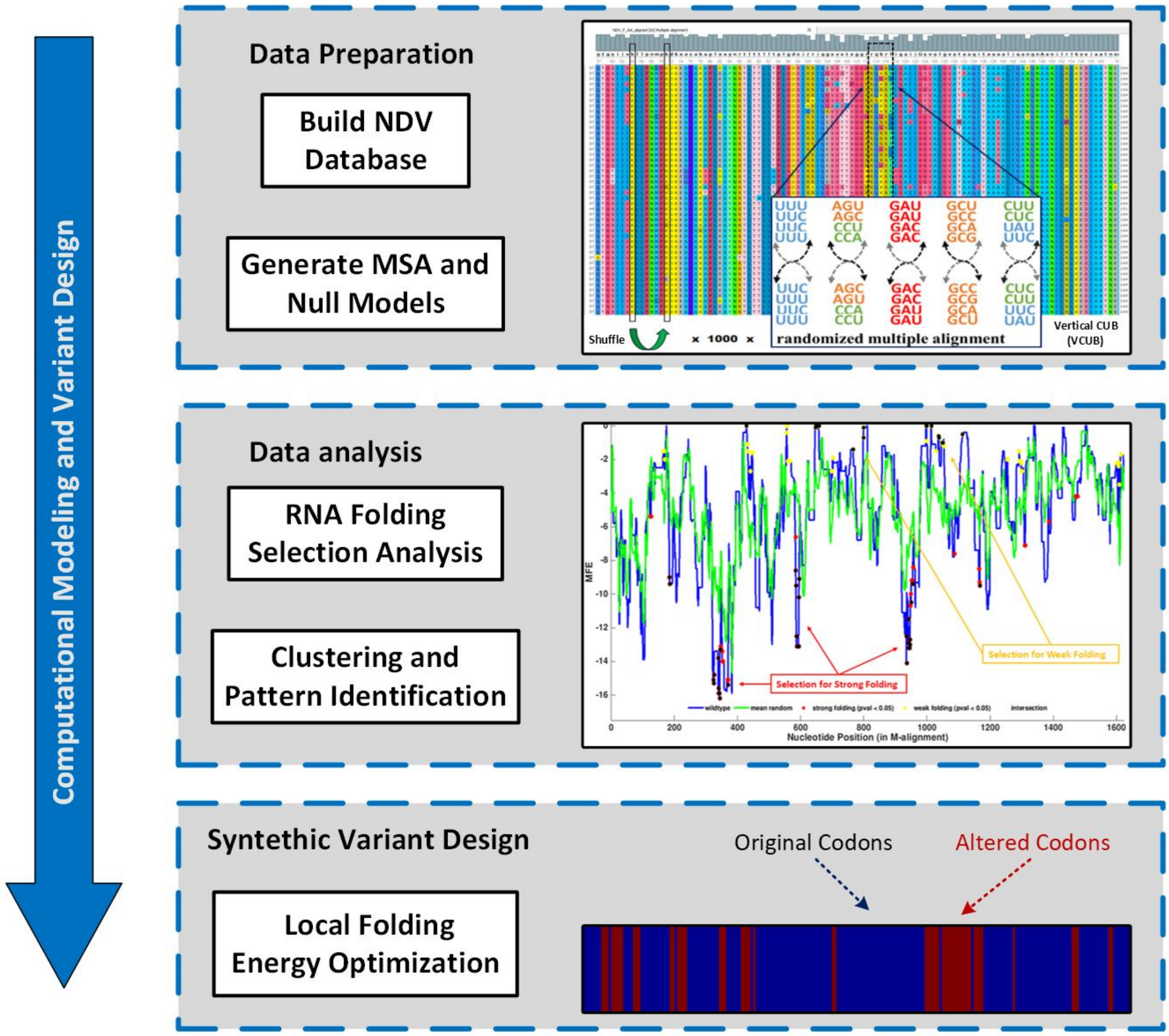


**Figure SF1: Scheme of the computational pipeline used for the rational design of attenuated synthetic NDV.** Top) Collection of NDV sequences, focusing on the fusion (F) protein, construction of a multiple sequence analysis (MSA) scheme, and generation of Null models of randomized sequences. Middle) RNA secondary structure (and folding) selection analysis, clustering, and pattern discovery. Bottom) Rational design of synthetic variants based on the folding selection patterns. The original codons are shown in blue while the altered synonymous codons are in marked red.

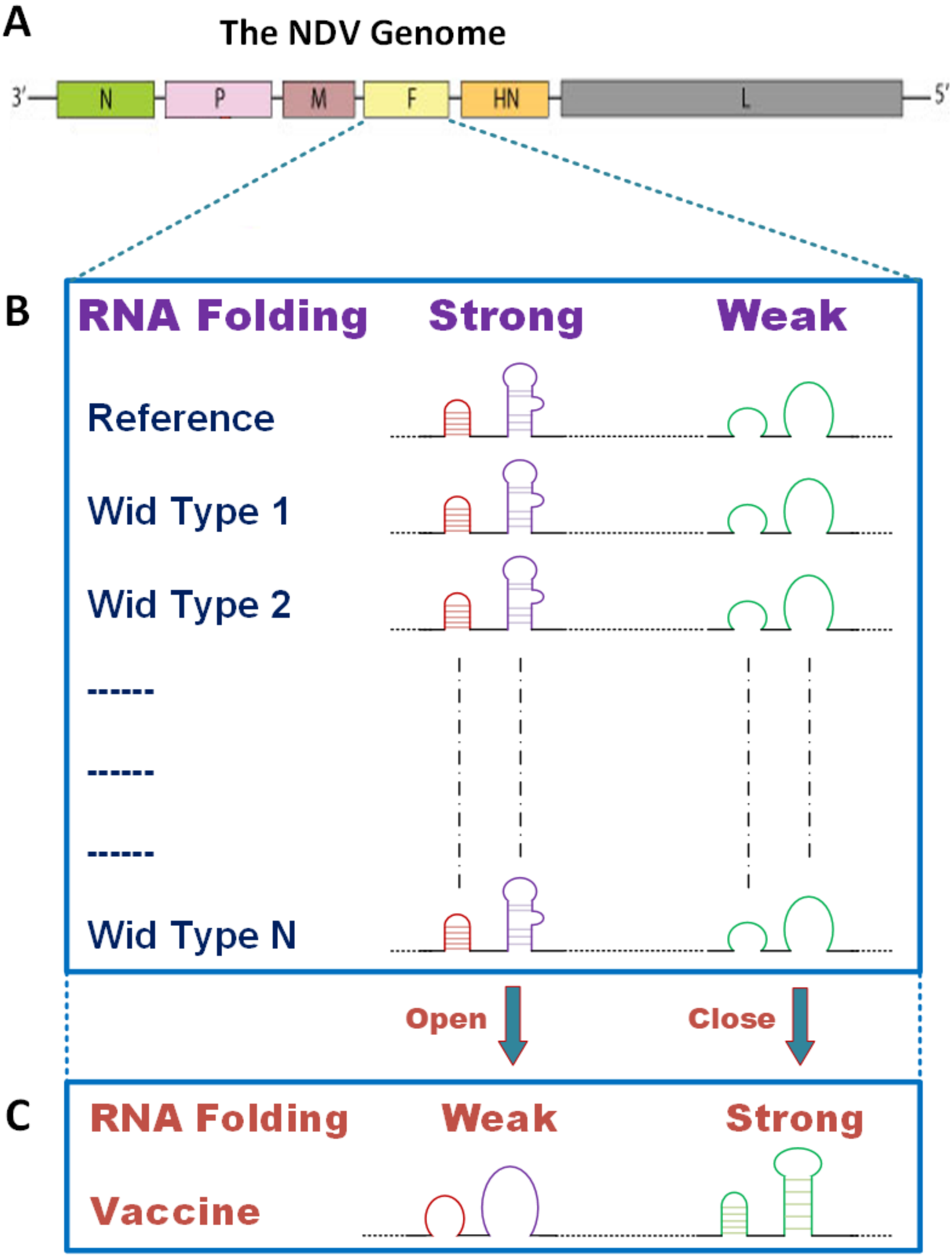


**Figure SF2: Visual abstract of the project.** A) The genome of the Newcastle Disease Virus (NDV) is composed of six main proteins: Nucleoprotein (N), Phosphoprotein (P), Matrix (M), Fusion (F) protein, Hemagglutinin-Neuraminidase (HN), and RNA polymerase (L). B) The reference and wildtype (WT) sequences of the F protein share some clustered sites with conserved RNA secondary structures for strong or weak folding *(i.e.*, closed or opened structures, respectively). C) Those sites were modified from strong to weak (or weak to strong), to generate an attenuated version of the virus.

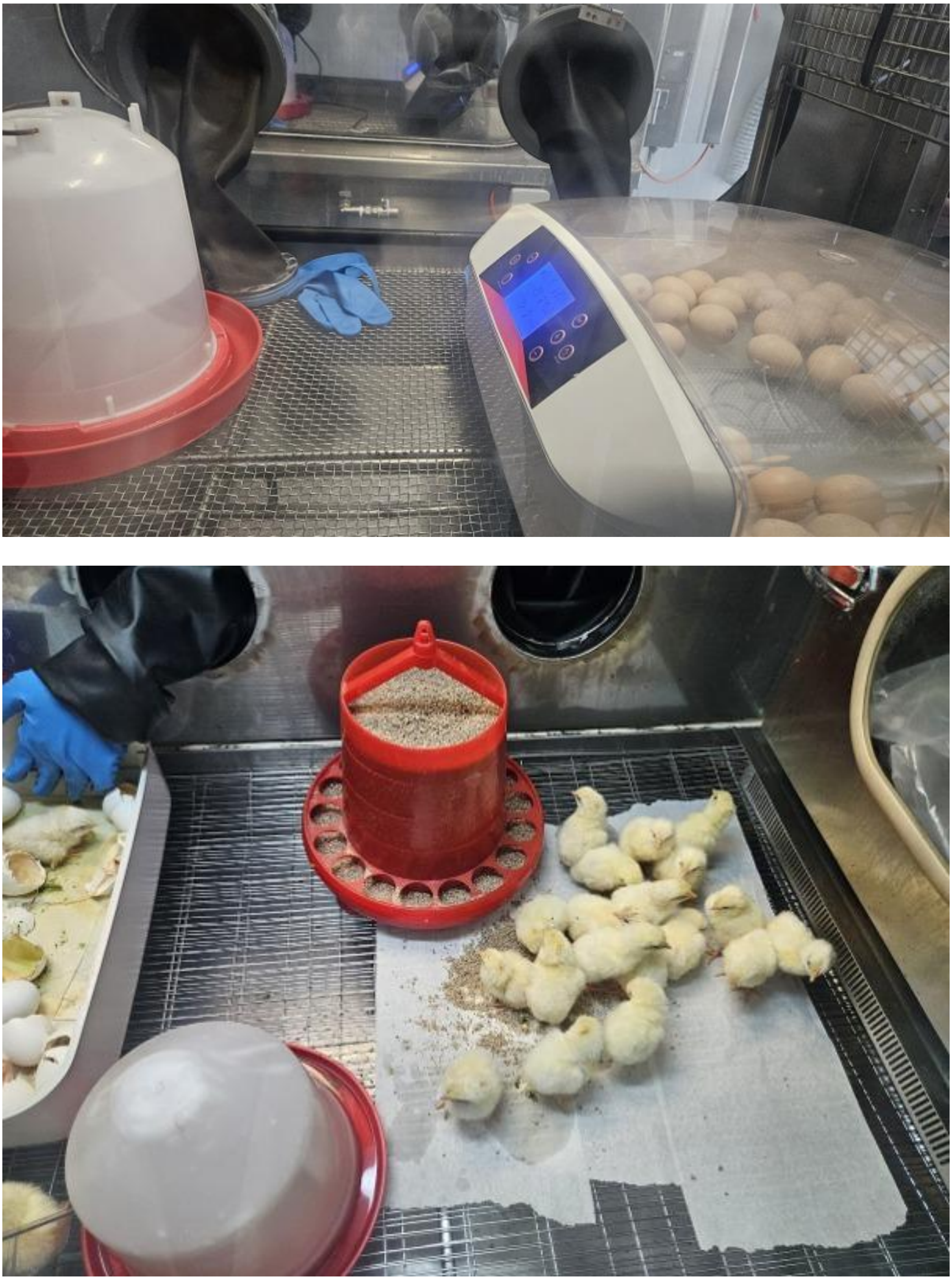

**Figure SF3. Incubation (top) and hatching (bottom) pictures.**

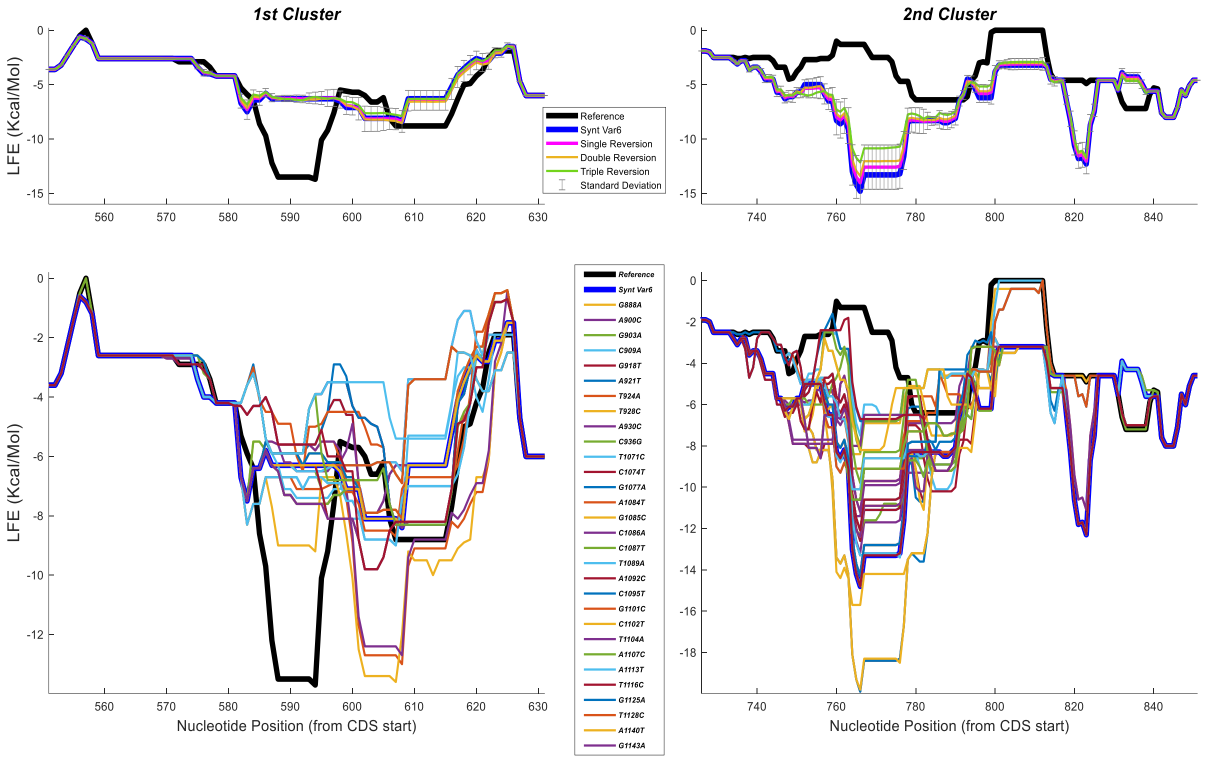


**Figure SF4. Reversion analysis.** We show the folding energy profiles for a single mutation, two adjacent mutation and three mutations in random locations, for the first and second selection clusters (left and right, respectively). Top) Average LFE profiles; standard deviation is shown for the single reversion. Bottom) single mutation profiles for each mutation introduced to the synthetic variant SynVar6.

### Supplemental Movies

**Movie SM1. Hatching movie No' 1.**

**Movie SM2. Hatching movie No' 2.**

### Supplemental Tables

**Table ST1. : Hatchability Rates**

**Table ST2. Serological monitoring (HI titers)**

**Table ST3. Challenge results (Protection percentage by group)**

### Appendix: Synthetic Variant Sequences

**Reference Sequence:** pNDVH_v2020, Newcastle disease virus isolate La Sota C5, complete genome, location: 4544...5743.

ACCGCGTGCCTGCGGAGCGTTAAAATCATTATCCAAGCAGGTACCCAACGCGCTGTCGCAGTGACCGCCGAC
CACGAGGTTACCTCTACTAAGCTGGAGAAGGGGCACACCCTTGCCAAATACAATCCTTTTAAGAAATAAGCTGC
GTCTCTGAGATTGCGCTCCGCCCACTCACCCAGATCATCATGACACAAAAAACTAATCTGTCTTGATTATTTACAG
TTAGTTTACCTGTCTATCAAGTTAGAAAAAACACGGGTAGAAGATTCTGGATCCCGGTTGGCGCCCTCCAGGTGC
AAGATGGGCTCCAGACCTTCTACCAAGAACCCAGCACCTATGATGCTGACTATCCGGGTTGCGCTGGTACTGAG
TTGCATCTGTCCGGCAAACTCCATTGATGGCAGGCCTCTTGCAGCTGCAGGAATTGTGGTTACAGGAGACAAAG
CCGTCAACATATACACCTCATCCCAGACAGGATCAATCATAGTTAAGCTCCTCCCGAATCTGCCCAAGGATAAGG
AGGCATGTGCGAAAGCCCCCTTGGATGCATACAACAGGACATTGACCACTTTGCTCACCCCCCTTGGTGACTCT
ATCCGTAGGATACAAGAGTCTGTGACTACATCTGGAGGGGGGAGACAGGGGCGCCTTATAGGCGCCATTATTGG
CGGTGTGGCTCTTGGGGTTGCAACTGCCGCACAAATAACAGCGGCCGCAGCTCTGATACAAGCCAAACAAAAT
GCTGCCAACATCCTCCGACTTAAAGAGAGCATTGCCGCAACCAATGAGGCTGTGCATGAGGTCACTGACGGAT
TATCGCAACTAGCAGTGGCAGTTGGGAAGATGCAGCAGTTTGTTAATGACCAATTTAATAAAACAGCTCAGGAATT
AGACTGCATCAAAATTGCACAGCAAGTTGGTGTAGAGCTCAACCTGTACCTAACCGAATTGACTACAGTATTCGG
ACCACAAATCACTTCACCTGCTTTAAACAAGCTGACTATTCAGGCACTTTACAATCTAGCTGGTGGAAATATGGATT
ACTTATTGACTAAGTTAGGTGTAGGGAACAATCAACTCAGCTCATTAATCGGTAGCGGCTTAATCACCGGTAACCC
TATTCTATACGACTCACAGACTCAACTCTTGGGTATACAGGTAACTCTACCTTCAGTCGGGAACCTAAATAATATGC
GTGCC

**Synthetic Variant No. 6:** based on the folding selection analysis (FOLD30_25_30; change ratio of 30%, with respect to the original LFE; 25 codons changed; 30 nucleotides changed).

ACCGCGTGCCTGCGGAGCGTTAAAATCATTATCCAAGCAGGTACCCAACGCGCTGTCGCAGTGACCGCCGAC
CACGAGGTTACCTCTACTAAGCTGGAGAAGGGGCACACCCTTGCCAAATACAATCCTTTTAAGAAATAAGCTGC
GTCTCTGAGATTGCGCTCCGCCCACTCACCCAGATCATCATGACACAAAAAACTAATCTGTCTTGATTATTTACAG
TTAGTTTACCTGTCTATCAAGTTAGAAAAAACACGGGTAGAAGATTCTGGATCCCGGTTGGCGCCCTCCAGGTGC
AAGATGGGCTCCAGACCTTCTACCAAGAACCCAGCACCTATGATGCTGACTATCCGGGTTGCGCTGGTACTGAG
TTGCATCTGTCCGGCAAACTCCATTGATGGCAGGCCTCTTGCAGCTGCAGGAATTGTGGTTACAGGAGACAAAG
CCGTCAACATATACACCTCATCCCAGACAGGATCAATCATAGTTAAGCTCCTCCCGAATCTGCCCAAGGATAAGG
AGGCATGTGCGAAAGCCCCCTTGGATGCATACAACAGGACATTGACCACTTTGCTCACCCCCCTTGGTGACTCT
ATCCGTAGGATACAAGAGTCTGTGACTACATCTGGAGGGGGGAGACAGGGGCGCCTTATAGGCGCCATTATTGG
CGGTGTGGCTCTTGGGGTTGCAACTGCCGCACAAATAACAGCGGCCGCAGCTCTGATACAAGCCAAACAAAAT
GCTGCCAACATCCTCCGACTTAAAGAGAGCATTGCCGCAACCAATGAGGCTGTGCATGAGGTCACTGACGGAT
TATCGCAACTAGCAGTGGCAGTTGGGAAGATGCAGCAGTTTGTTAATGACCAATTTAATAAAACAGCTCAGGAgTT
AGACTGCATaAAgATTGCcCAGCAAGTgGGaGTtGAGtTaAACCTcTACCTAACCGAATTGACTACAGTATTCGGAC
CACAAATCACTTCACCTGCTTTAAACAAGCTGACTATTCAGGCACTTTACAATCTAGCTGGTGGAAATATGGATTAC
TTATTGACTAAGTTAGGTGTAGGGAAtAAcCAgCTCAGCagccTtATaGGcAGCGGgcTtATaACCGGaAAtCCTATTC
TgTAtGACTCACAGACaCAgCTCTTGGGTATACAGGTAACTCTACCTTCAGTCGGGAACCTAAATAATATGCGTGCC

Note: changed nucleotides are marked in lowercase letters.
